# Novel Biomarkers and Distinct Transcriptomic Profile of Barrett’s Esophagus Epithelial Stem Cells

**DOI:** 10.1101/2023.08.07.552218

**Authors:** Katie L. Alexander, Lesley E. Smythies, Kondal R. Kyanam-Kabir-Baig, Emily Poovey, David K. Crossman, Phillip D. Smith, Shajan Peter

**Affiliations:** Department of Medicine (Gastroenterology), University of Alabama at Birmingham, Birmingham, AL; Department of Genetics, University of Alabama at Birmingham, Birmingham, AL

**Keywords:** Esophageal Adenocarcinoma, Barrett’s Esophagus, Organoids

## Abstract

Barrett’s esophagus, a metaplastic condition that originates in the distal esophagus, is the only known precursor lesion for the development of esophageal adenocarcinoma, which has a devasting 5-year survival rate of <20%. The large number of subjects diagnosed with Barrett’s esophagus, and therefore at higher risk for esophageal adenocarcinoma, underscores the necessity for biomarkers that would benefit surveillance and potentially early treatment. To address this, we generated epithelial stem cell organoids from normal gastric cardia, non-dysplastic and dysplastic Barrett’s esophagus, and esophageal and gastric adenocarcinoma. Interestingly, non-dysplastic and dysplastic Barrett’s esophagus displayed higher expression of multiple archetypical cancer-associated genes compared with both esophageal and gastric adenocarcinoma in addition to expression of the novel biomarker CT83. ST6GAL1, a Golgi sialyltransferase upregulated in multiple epithelioid cancers, was strongly upregulated in dysplastic Barrett’s esophagus at both mRNA and protein levels. ST6GAL1 protein also was highly expressed in esophageal adenocarcinoma, suggesting that regulation of ST6GAL1 may play a role in Barrett’s esophagus progression to esophageal adenocarcinoma and serve as a potential biomarker of the development of esophageal cancer.

## Introduction

Esophageal cancer has a devasting 5-year survival rate of <20%(Thrift, 2021) and is composed of two histological subtypes, esophageal squamous cell carcinoma (ESCC) and esophageal adenocarcinoma (EAC), the latter rapidly increasing among Western populations during the past 40 years(Thrift, 2021). Barrett’s esophagus (BE), a metaplastic condition that originates in the distal esophagus, is characterized by the replacement of squamous epithelium with columnar epithelium with gastric and intestinal features and is the only known precursor lesion for the development of EAC(Rustgi & El-Serag, 2014). BE occurs in up to 15% of subjects with gastroesophageal reflux disease (GERD) and is estimated to affect 0.5-2.0% of the general population(Schmidt *et al*, 2022). Although metaplasia in BE advances to low-grade dysplasia (LGD) or high-grade dysplasia (HGD) in only a subset of patients, such progression significantly augments the risk for EAC(Iyer & Chak, 2023; Killcoyne & Fitzgerald, 2021). Thus, the large number of subjects diagnosed with BE, consequently at higher risk for EAC, underscores the importance of identifying biomarkers of disease progression that would inform surveillance and potentially enable earlier diagnosis and treatment.

The cell of origin for BE metaplasia, and whether it is derived from the esophagus or stomach, has been the subject of intense debate(Jiang *et al*, 2017; Nowicki-Osuch *et al*, 2023; Nowicki-Osuch *et al*, 2021; Owen *et al*, 2018; Que *et al*, 2019; Rhee & Wang, 2018). Previous work from Owen *et al* demonstrated that Barrett’s esophagus had more transcriptional overlap with esophageal submucosal gland (SMG) cells compared with gastric or duodenal cells(Owen *et al*., 2018), suggesting BE is derived from the SMG cells of the esophagus. In contrast, Nowicki-Osuch *et al* recently showed that ectopic expression of c-MYC and HNF4A in normal gastric cardia epithelial cells resulted in the expression of BE-associated genes(Nowicki-Osuch *et al*., 2021), suggesting BE most likely arises from gastric cardia cells(Nowicki-Osuch *et al*., 2021). In agreement with the latter findings, data from The Cancer Genome Atlas Research Network suggests that EAC is more similar to chromosomally unstable GAC than ESCC, which more closely resembles squamous cancers at other sites than EAC(2017). Further, individual epithelial cells from both BE intestinal metaplasia and gastric intestinal metaplasia were shown recently to share a phenotype, with both cell populations able to simultaneously express gastric and intestinal features at the single cell level(Nowicki-Osuch *et al*., 2023). These findings together support the argument that the cell of origin for intestinal metaplasia in BE arises from the gastric cardia. In this study, we compare the transcriptional profile of epithelial stem cells, the progenitors of the epithelium, derived from non-dysplastic (ND BE) and dysplastic (LGD and HGD) BE with epithelial stem cells from the normal gastric cardia to identify potential novel biomarkers of BE, which may benefit early BE diagnosis, which heavily relies on histopathology. Further, we probe epithelial stem cells from EAC to investigate potential biomarkers of BE disease progression that would enhance surveillance and potentially lead to early treatment of EAC.

## Results

In the setting of BE, the stratified squamous epithelium is replaced by single-layered columnar epithelium with intestinal features that may include goblet cells. To investigate BE epithelial stem cells, the progenitors of BE metaplastic epithelium, for potential novel biomarkers of BE and disease progression, we generated epithelial stem cell organoids from biopsies of healthy gastric cardia (control) and esophagus from subjects with BE or EAC (**Fig. 1A**). We first examined organoids generated from normal cardia, BE, and EAC tissue for expression of *LGR5*, a conical stem cell marker(Barker *et al*, 2010; Barker *et al*, 2007); *SOX9*, a transcriptional factor associated with BE(Wang *et al*, 2010) and multiple cancers(Aguilar-Medina *et al*, 2019); and *CDX2*, an intestinal epithelial cell transcriptional factor associated with metaplasia in BE(Chen *et al*, 2011) (**Fig. 1B**). Expression of *LGR5* and *SOX9* were similar across all samples, but BE-derived organoids highly expressed *CDX2* compared with normal gastric cardia-derived and EAC-derived organoids (**Fig. 1B**), consistent with the important role of CDX2 in intestinal epithelial cell differentiation.

**Figure 1-.**
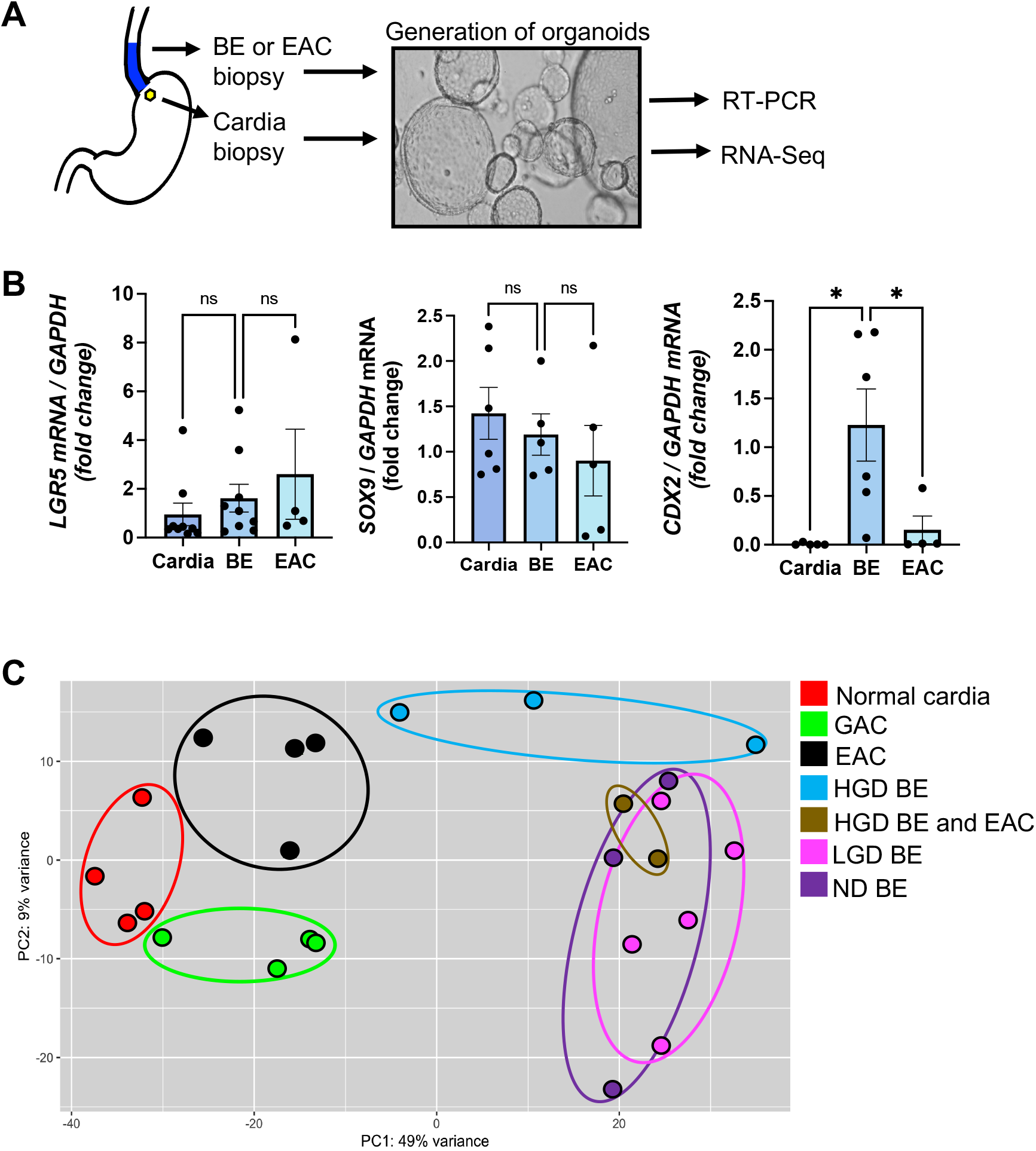
Transcriptional diversity of epithelial organoids from BE, EAC and GAC and normal gastric cardia. **A** Experiment sequence schematic. **B** Human epithelial organoids were generated from normal cardia (n=5-9), BE (n=5-9) or EAC (n=4-5) biopsies and analyzed for *LGR5*, *SOX9* and *CDX2* mRNA expression by real time PCR. (Data are shown as mean +/− SEM, one-way ANOVA, significance: *, *p* <0.05) **C** β-diversity of mRNA gene expression in organoids generated from ND BE (n=3, purple), LGD BE (n=5, pink), HGD BE (n=3, blue), HDG BE and EAC (n=2, brown), EAC (n=4, black), GAC (n=4, green) and normal gastric cardia (n=4, red) determined by RNA-Seq and displayed as a PCA plot with each dot representing a single subject.

To gain insight into the breath of gene expression in epithelial stem cells at each stage of BE progression, we next performed comparative transcriptomic analysis of epithelial stem cell organoids generated from non-dysplastic BE (ND BE), low-grade dysplasia BE (LGD BE), high-grade dysplasia BE (HGD BE), BE with HGD and EAC (HGD BE and EAC) and EAC (**Fig. 1C**). Given the evidence that BE may originate from gastric cardia cells(Nowicki-Osuch *et al*., 2023; Nowicki-Osuch *et al*., 2021), we included in this analysis epithelial stem cells generated from normal gastric cardia and gastric adenocarcinoma (GAC) (**Fig. 1C**). Principal Component Analysis (PCA) of genomic variation among the groups of epithelial stem cells showed that ND BE and LGD BE overlapped transcriptomically and that HGD BE was transcriptomically intermediate between ND BE / LGD BE and EAC (**Fig. 1C**). The transcriptome profile of EAC stem cells closely clustered with GAC stem cell profiles (**Fig. 1C**), suggesting commonality, as previously reported(2017; Nowicki-Osuch *et al*., 2023).

To further elucidate the transcriptional evolution of epithelial stem cells from patients at each stage of BE progression (**Fig. 1C**), we profiled organoids generated from ND BE, LGD BE, HGD BE for potential biomarker, immune response, and cancer-associated genes. We compared the gene expression of each BE group, in addition to gene expression of EAC and GAC organoids, to normal cardia-derived organoid gene expression (**Fig. 2**), given that that gastric cardia is the most likely source of metaplastic BE cells(Nowicki-Osuch *et al*., 2023; Nowicki-Osuch *et al*., 2021; Peters *et al*, 2019). The homeobox transcription factors *CDX2,* which drives glandular cell fate, as well as *HOXA13, HOXB3* and *HOXC10*, which regulate cell differentiation of oncogenic potential, and *PTGS2,* which drives prostaglandin biosynthesis, were strongly upregulated in both non-dysplastic and dysplastic BE but not in EAC (**Fig. 2**), confirming a classical BE signature(Killcoyne & Fitzgerald, 2021). Strong upregulation of *PTGS2* in BE-derived organoids was recapitulated in a second cohort of BE samples (**Fig. EV1a**), further corroborating a BE signature. Notably, we detected increased expression of the mucin genes *MUC13* and *MUC7* in both ND BE and dysplastic BE (**Fig.2**). In addition, the expression of 7 of the 13 immune regulation genes analyzed were elevated in each BE group (**Fig. 2**), consistent with the well-recognized immune function of mucosal epithelial cells(Peterson & Artis, 2014). Non-dysplastic and dysplastic BE epithelial stem cells displayed more pronounced upregulation of 18 of the 34 stemness- and cancer-associated genes analyzed, including *CEACAM5*, *CEACAM6*, *RUNX2, RUNX3* and *PAX9,* than either EAC or GAC (**Fig. 2**). Strikingly, cancer testis antigen 83 (*CT83)*, a gene not previously linked to BE and not typically expressed in normal tissue outside testis(Chen *et al*, 2021), showed remarkable specificity for BE epithelial stem cells compared with EAC and GAC (**Fig. 2**). We further confirmed elevated *CT83* expression and specificity in BE with an additional cohort of BE- and EAC-derived stem cell organoids (**Fig. EV1b**).

**Figure 2-.**
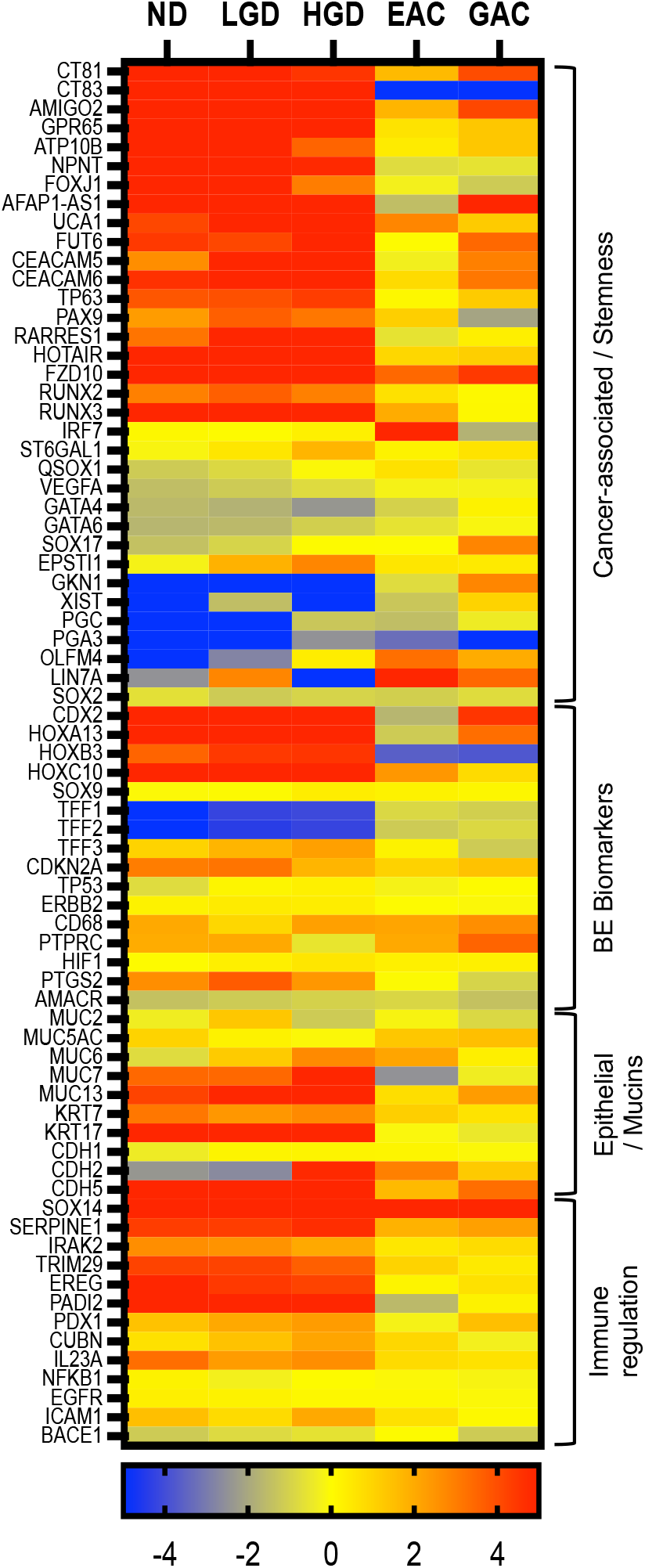
Epithelial stem cells from BE, EAC and GAC have distinct transcriptomic profiles. Heat map displaying epithelial stem cell mRNA gene expression in organoids generated from ND BE (n=3), LGD BE (n=5), HGD BE (n=3), EAC (n=4), GAC (n=4) compared with the mRNA expression of the indicated genes in normal gastric cardia-derived organoids (n=4, control). (Scale: log base 2-fold change, range −5 to 5)

A key feature of malignant potential is dysregulated apoptosis. In this connection, we have reported that glycosyltransferase ST6GAL1, which catalyzes the addition of α2,6-linked sialic acid to TNFR1 and thus disrupts TNF-mediated homeostatic apoptosis, is upregulated in gastric adenocarcinoma epithelial cells(Alexander *et al*, 2020). ST6GAL1 is also upregulated in multiple other cancers(Alexander *et al*., 2020; Britain *et al*, 2017; Chakraborty *et al*, 2018; Dorsett *et al*, 2021), including esophageal and gastric adenocarcinoma(Chandrashekar *et al*, 2017; Chandrashekar *et al*, 2022) (**Fig. EV2a**). Transcriptomic analysis revealed an upregulation of *ST6GAL1* mRNA in LGD BE, and to a greater extent in HGD BE, compared with normal cardia (**Fig. 2**). We confirmed *ST6GAL1* mRNA was significantly upregulated in organoids derived from dysplastic (LGD and HGD) BE (**Fig. 3A**) in an additional cohort but intriguingly did not observe an increase in EAC-derived organoids (**Fig. 3A**). Further, *ST6GAL1* expression was retained in terminally differentiated BE epithelial cells generated from both non-dysplastic and dysplastic BE-derived stem cell organoids, similar to epithelial cells in gastric adenocarcinoma, as we have previously reported(Alexander *et al*., 2020). However, *ST6GAL1* was not present at high levels in monolayers generated from normal cardia (**Fig. 3B**), as we have shown to be present in normal gastric antrum(Alexander *et al*., 2020), or EAC-derived organoids (**Fig. 3B**). The unexpected lack of upregulated of ST6GAL1 expression in EAC-derived organoids and monolayers (**Fig. 3A,B**) is in contrast to public data from The Cancer Genome Atlas (TCGA), which we analyzed using The University of Alabama at Birmingham Cancer data analysis portal (UALCAN)(Chandrashekar *et al*., 2017; Chandrashekar *et al*., 2022). The UALCAN database(Chandrashekar *et al*., 2017; Chandrashekar *et al*., 2022) revealed a significant upregulation of *ST6GAL1* in EAC compared with esophageal squamous cell carcinoma and healthy controls (**Fig. EV2b**). Further, high *ST6GAL1* expression is negatively associated with esophageal cancer patient survival (**Fig. EV2c**), similar to ovarian carcinoma(Garnham *et al*, 2019; Jones *et al*, 2023; Wichert *et al*, 2018). One possible explanation for the differences seen between the TCGA database and our data is that the TCGA database compares normal and tumorigenic tissue, and normal esophageal squamous epithelium does express appreciable levels of ST6GAL1 (**Fig. EV3**). In contrast, we analyzed ST6GAL1 expression in columnar epithelial cells in gastric cardia, BE and EAC. Additionally, the level of active ST6GAL1 maybe be similar in BE and EAC. ST6GAL1 was recently demonstrated to be post-transcriptionally regulated by the disulfide catalyst QSOX1(Ilani *et al*, 2023), which activates ST6GAL1 through disulfide bond formation(Ilani *et al*., 2023). QSOX1 also has been shown to play a role in protein folding, cell cycle regulation, extracellular matrix remolding(Portes *et al*, 2008), and cancer detection and prognosis(Antwi *et al*, 2009; Katchman *et al*, 2013; Lake & Faigel, 2014). To this end, we measured *QSOX1* expression in BE and EAC organoids and differentiated monolayers and demonstrated similar expression levels at the mRNA level (**Fig. 4A**) and prominent QSOX1 protein expression in both BE and EAC (**Fig. 4B**).

**Figure 3-.**
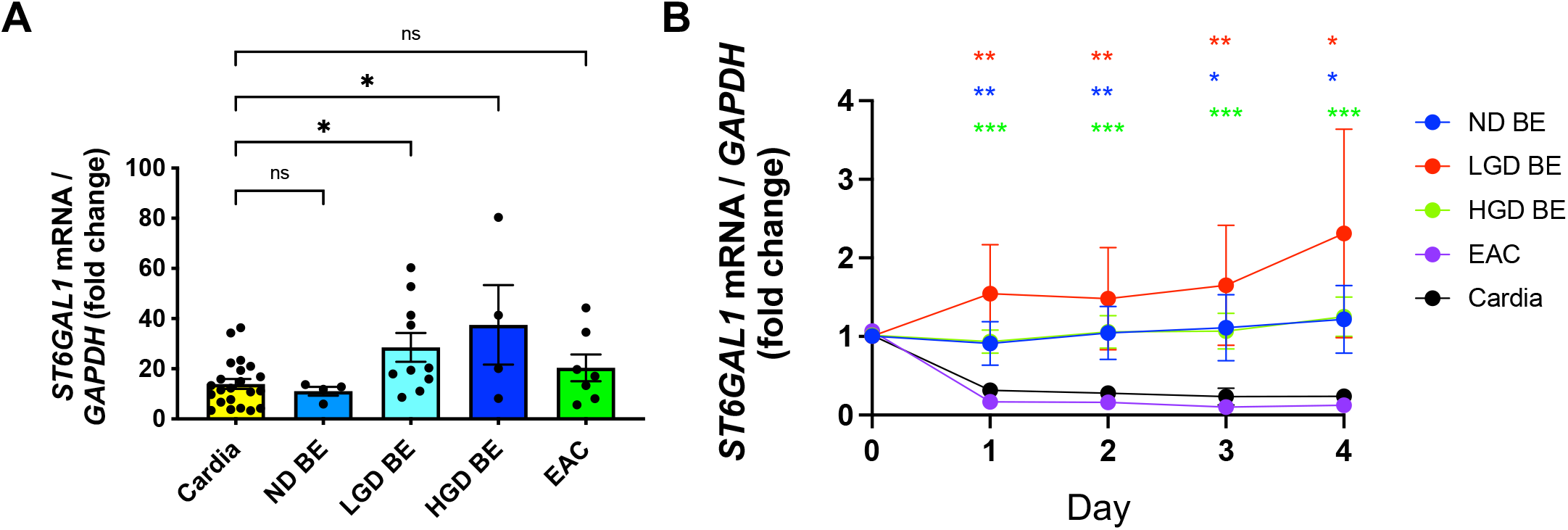
ST6GAL1 is highly expressed in dysplastic BE. **A** Epithelial organoids generated from biopsies from normal gastric cardia (n=22, control), ND BD (n=4), LGD BE (n=10), HGD BE (n=4) or EAC (n=7) were analyzed for *ST6GAL1* gene expression by real time PCR. (Data are shown as mean +/− SEM, one-way ANOVA, significance: *, p<0.05). **B** Epithelial cells monolayers were generated from ND BE (n=5), LGD BE (n=3), HGD BE (n=4), EAC (n=2) and cardia (n=6, control) organoids as previously described(Alexander *et al*., 2020). Fold change in *ST6GAL1* mRNA was measured on day 0 (organoids) and days 1-4 (monolayer) by real time PCR. Data are shown as mean +/− SEM, unpaired t test between each group and control for each day, significance: *, *p* <0.05, **, p <0.005, ***, p <0.0001).

**Figure 4-.**
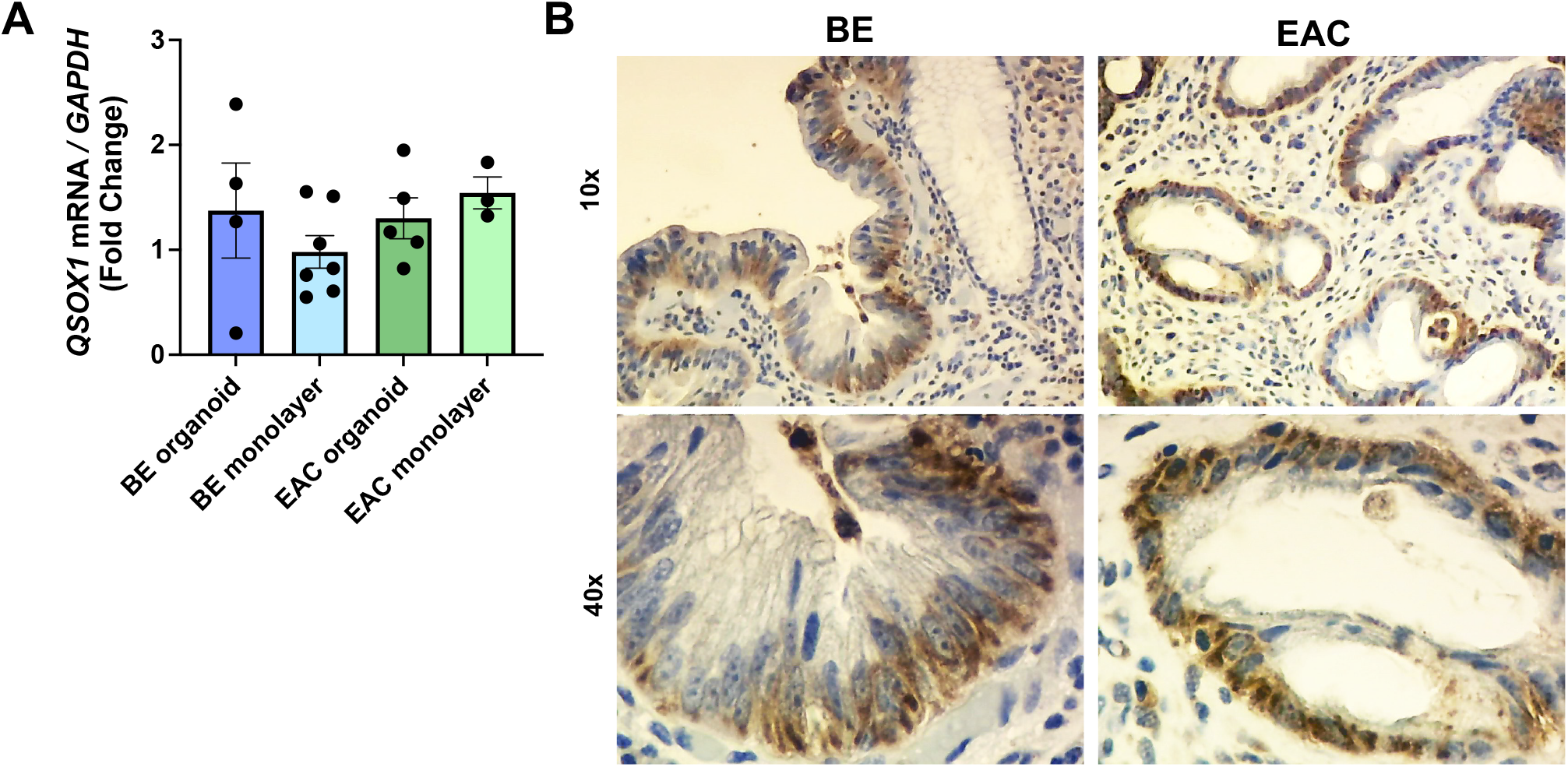
QSOX1 is highly expressed in BE and EAC. **A** Epithelial organoids and organoid-derived monolayers were generated from BE (n=4-7) or EAC (n=3-5) tissue and analyzed for *QSOX1* gene expression by real time PCR. (Data shown as mean +/− SEM). **B** Esophageal tissue from subjects with BE (ND BE shown) or EAC was stained for QSOX1 (DAB) by immunohistochemistry (n=3 each, representative images shown, 10× and 40×).

To determine the level of transcribed ST6GAL1 in BE and EAC, we next examined ST6GAL1 protein expression in BE and EAC. Both organoids and organoid-derived differentiated epithelial cell monolayers derived from subjects with BE expressed high levels of ST6GAL1 protein (**Fig. 5A,B**). However, in contrast to the low mRNA expression of ST6GAL1 in EAC, levels of ST6GAL1 protein were also elevated in EAC-derived organoids and organoid-derived differentiated epithelial cell monolayers (**Fig. 5A,B**). These data are in agreement with our previous work that showed that ST6GAL1 protein expression increases progressively during gastric premalignancy and is expressed at the highest level in gastric adenocarcinoma(Alexander *et al*., 2020). Since patient biopsies are used to confirm a diagnosis of BE, we next examined tissue biopsies from subjects with BE and EAC. High ST6GAL1 protein expression also was present in the epithelial cells (**Fig. 5C**), positioning ST6GAL1 as a potential biomarker for BE that may progress to EAC.

**Figure 5-.**
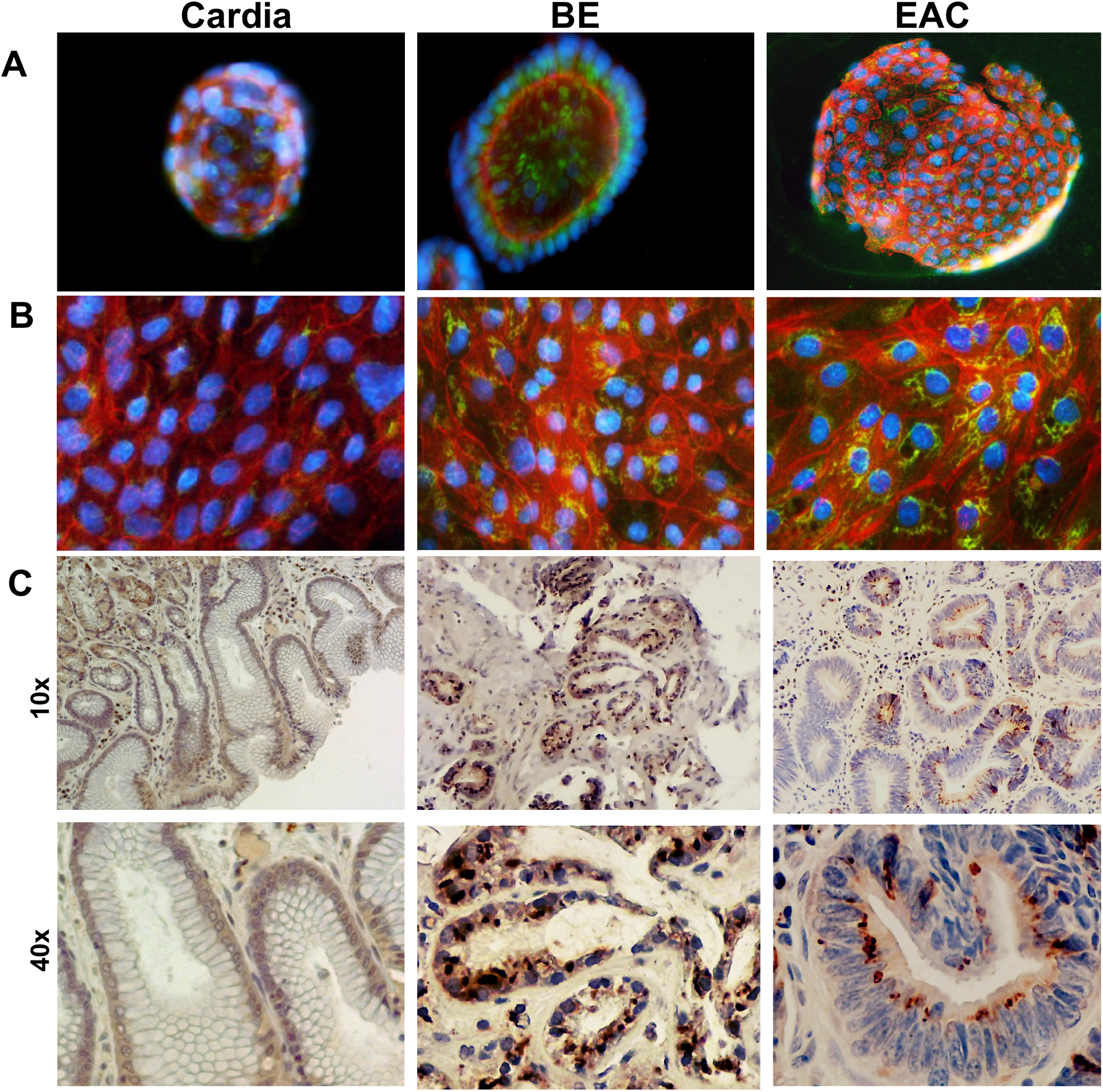
ST6GAL1 protein is highly expressed in BE and EAC organoids, organoid-derived monolayers, and tissue. **A,B (A)** Cardia, BE-(LGD shown) or EAC-derived organoids or **(B)** day 2 organoid-derived monolayers stained with antibodies for ST6GAL1 (FITC), phalloidin (Alexa Fluor 594), and DAPI (*n* = 3 each, representative donors shown, 20x). **C** Cardia tissue from healthy donors or esophageal tissue from subjects with BE (LGD shown) or EAC stained with antibodies to ST6GAL1 (DAB) by immunohistochemistry (n=3, representative images shown, 10× and 40×).

## Discussion

Barrett’s esophagus is the only known precursor lesion for the development of EAC; however, the majority of subjects with BE will not progress to EAC and not all subjects with EAC have evidence of BE(Sawas *et al*, 2018). Current classification of BE is based largely on histology, which can vary depending on pathologist-specific scoring or physician-specific sampling; therefore, a more definitive diagnostic methodology is desirable. Nowicki-Osuch and colleagues(Nowicki-Osuch *et al*., 2021) recently showed that undifferentiated BE cells, regardless of the presence of metaplastic precursors, expressed markers present in EAC(Nowicki-Osuch *et al*., 2021), underscoring the potential importance for identifying markers of BE diagnosis beyond histopathology. Consequently, a biomarker or combination of biomarkers that identify subjects with BE at high-risk for progressing to EAC would inform surveillance and early diagnosis and potentially enhance successful intervention. To this end, recent strides in the detection of BE have included Cytosponge-TFF3(Fitzgerald *et al*, 2020), a non-invasive detection method reported to increase detection of BE by measuring the expression of *TFF3* in the distal esophagus(Fitzgerald *et al*., 2020). In addition, TissueCypher® technology provides a risk score for the development of HGD or EAC within 5 years of sampling based on the expression of *CDKN2A, TP53, ERBB2, CD68, PTPRC, HIF1, PTGS2, AMACR, KRT20*(Diehl *et al*, 2021), rather than histology or BE segment length alone. Of note, we report the gene expression level of each of these biomarkers in epithelial stem cells from BE, EAC and GAC, with the exception of *KRT20,* which was not detected in our analysis. Implementation of TissueCypher® into clinical practice has been reported to impact surveillance intervals and physician decision-making(Diehl *et al*., 2021) but depends on an initial diagnosis of BE and has not shown statistical significance in identifying high-risk BE.

Our findings uncover distinct transcriptome profiles in epithelial stem cells derived from the esophageal tissue of subjects with BE and EAC. Interestingly, ND BE and dysplastic BE stem cells largely show similar transcriptomes, with some caveats. *CT8*3 mRNA expression had a strong specificity for BE-derived epithelial stem cells compared with EAC and GAC. Intriguingly, while not expressed in EAC, *CT83* has recently been reported to be overexpressed in several cancers and possesses notable specificity for triple-negative breast cancer(Chen *et al*., 2021). The high specificity of *CT83* in BE-derived organoids suggests a potential novel epithelial biomarker of BE. While the stark differences in CT83 expression in BE and EAC are intriguing, further studies utilizing a larger cohort of BE subjects consisting of progressors and non-progressors is necessary to determine whether the low expression of CT83 in EAC suggests that it is indicative of low-risk BE.

*ST6GAL1* was significantly upregulated in organoids derived from dysplastic (LGD and HGD) BE but not ND BE compared with cardia-derived organoids. While we did not see an increase in *ST6GAL1* in EAC at the mRNA level, we demonstrated elevated ST6GAL1 expression in both BE and EAC at the protein level. This discrepancy may be due to regulation of ST6GAL1 activity. The disulfide catalyst QSOX1 has recently been shown to regulate ST6GAL1 activity through disulfide bond formation(Ilani *et al*., 2023). Interestingly, we detected no significant difference in *QSOX1* mRNA expression between BE- and EAC-derived organoids and organoid-derived monolayers, suggesting *QSOX1* may be a factor in the regulation of ST6GAL1 activity and protein expression. In addition to ST6GAL1, non-dysplastic and dysplastic BE epithelial stem cells displayed more pronounced upregulation of 18 of the 34 stemness- and cancer-associated genes analyzed, including *CEACAM5*, *CEACAM6*, *RUNX2, RUNX3* and *PAX9,* than either EAC or GAC. Whether this upregulation is due to the dynamic nature of BE metaplasia or marks causative drivers of disease progression needs further investigation.

In summary, our study characterizes unique gene expression profiles in esophageal epithelial stem cells for each stage of BE leading to, and including, EAC. Further, we used comparative transcriptomic analysis to identify groups of genes that could serve as biomarkers of disease. Although cross-sectional, our findings enlarge the framework for future study of the genes that identify progressing and non-progressing BE and that could be manipulated using organoid models to determine their role in driving BE progression to EAC.

## Materials and Methods

### Human tissue samples

Biopsy specimens were obtained from the gastric cardia of healthy adults without mucosal pathology, BE segments (with or without dysplasia), and esophageal or gastric cancer from subjects with histologically confirmed pathology undergoing clinically indicated endoscopy at our institution.

### Stem cell organogenesis

Epithelial stem cell organoids were generated from tissue specimens in the UAB Stem Cell Organogenesis Unit. Briefly, biopsies were finely minced and digested for up to 50 min at 37°C in collagenase (2 mg/mL Collagenase Type 1; Corning), pipetting every 10 minutes. The collagenase then was neutralized with washing buffer (Advanced DMEM/F-12 (Sigma), 10% FBS (Atlanta Biologicals), L-glutamine 2 mM (Fisher Scientific), penicillin 100 U/mL and streptomycin 0.1 mg/mL (Fisher Scientific), gentamycin (50 μg/mL; Corning), fungizone (2.5 μg/mL; Fisher Scientific), and the cell suspension filtered through a 70-μm screen and centrifuged (200 g for 5 min at RT). Pelleted cells were either washed and pelleted again, or immediately suspended in Matrigel (Corning) in an ice-cold 24-well plate (15 μL/well; Corning). The Matrigel was allowed to polymerize (15 min, 37°C) with the plate upside-down to prevent the epithelial cells from attaching to the plate surface, after which culture medium (450 μL/well) containing a 50% mix of L-WRN fibroblast cell line conditioned media (CM) and primary culture medium (Advanced DMEM/F-12; Sigma) both supplemented with 20% fetal bovine serum (Atlanta Biologicals), L-glutamine 2 mM (Fisher Scientific), penicillin 100 U/mL and streptomycin 0.1 mg/mL (Fisher Scientific) was added. The CM was further supplemented with TGF-β R1 inhibitor (10 μM SB-431542, Selleck Chemicals), ROCK inhibitor (10 μM Y-27632; R&D Biosystems), gentamycin (50 μg/mL; Corning) and fungizone (2.5 μg/mL; Fisher Scientific) before addition to the wells. Organoids were maintained at 37°C and 5% CO_2_ and passaged every 4-7 days.

### Epithelial cell monolayers

Organoid cultures were dissociated in trypsin, washed, passed through a 40-μm screen (Corning) to remove cell clusters, resuspended in 5% L-WRN fibroblast cell line conditioned media in primary culture medium, counted and seeded onto an 8-well glass chamber slide (200 μL/well) or 48-well cell culture plate (350 μL/well; both Corning), which was previously coated with Matrigel (1:40 dilution in PBS). The individual stem cells were allowed to differentiate, reaching confluence in approximately 2 days.

### Quantitative PCR

Quantitative polymerase chain reaction (qPCR) was performed using a QuantStudio 3 (Thermo Fisher Scientific) instrument. Briefly, organoids or monolayers were lysed in RLT buffer (Qiagen), and RNA was extracted using the RNeasy kit (Qiagen). Equal amounts of RNA were used to generate cDNA with the High Capacity cDNA Reverse Transcription Kit (Fisher). qPCR experiments used TaqMan gene expression assay primers (Thermo Fisher) and TaqMan Fast Advanced Master Mix (Thermo Fisher) to generate reactions, and the comparative CT method was used to analyze the corresponding data.

### RNA-Seq

Epithelial stem cell organoids were lysed in RLT buffer (Qiagen), and RNA was extracted using the RNeasy kit (Qiagen). The extracted RNA was analyzed on the Illumina NextSeq 500 platform through the UAB Genomics Core. Raw sequence reads were aligned to the human genome from Gencode (GRch38 p13 Release 32) using STAR (version 2.7.4a; parameters used: --outReadsUnmapped Fastx --outSAMtype BAM SortedByCoordinate -- outSAMattributes All)(Dobin *et al*, 2013). Transcript abundances were then calculated from the STAR Alignments using HTSeq-count (version 0.11.1; parameters used: -m union -r pos -t exon -i gene_id -a 10 -s no -f bam)(Anders *et al*, 2014). Raw transcript abundances were then normalized and differential expression was calculated using DESeq2 following their vignette(Love *et al*, 2014).

### Statistical analysis

Student’s *t*-test or 1-way ANOVA followed by Turkey post-test was performed using Prism GraphPad software with p<0.05 considered statistically significant. Significance is indicated as *, p<0.05; **, p<0.01; ***, p<0.001; and ****, p<0.0001.

### Study approval

All human specimens were obtained after written informed consent approved by the UAB IRB and abide by the Declaration of Helsinki Principles.

## Abbreviations

BE: Barrett’s esophagus
LGD: low grade dysplasia
HGD: high grade dysplasia
NDBE: non-dysplastic Barrett’s esophagus
EAC: esophageal adenocarcinoma
GAC: gastric adenocarcinoma
GERD: gastro-esophageal reflux disease

## Funding Sources

This work was supporting by the University of Alabama at Birmingham (UAB) Health Services Foundation Grant Award, UAB Stem Cell Organogenesis Unit and National Institutes of Health Grants HD088954 and 2T32A1007051-41

## Author Contributions

**KLA:** Conceptualization: Equal, Formal Analysis: Lead, Investigation: Lead, Methodology: Equal, Data Curation: Lead, Writing-original draft: Lead, Writing-Review and Editing: Equal. Resources: Supporting **LES:** Methodology: Equal, Formal Analysis: Supporting, Writing-Review and Editing: Equal, Data Curation: Supporting. **KRKKB:** Writing-Review and Editing: Equal, Resources: Supporting. **DKC**: Formal Analysis: Supporting, Methodology: Supporting, Data Curation: Supporting **PDS:** Writing-Review and Editing: Equal, Data Curation: Supporting, Resources: Supporting. **SP:** Conceptualization: Equal, Writing-Review and Editing: Equal, Resources: Lead

Study materials will be made available upon request.

## Conflicts of Interests

The authors have no conflicts of interest to declare.

## Expanded View Figure Legends

**Expanded View Figure 1-.**
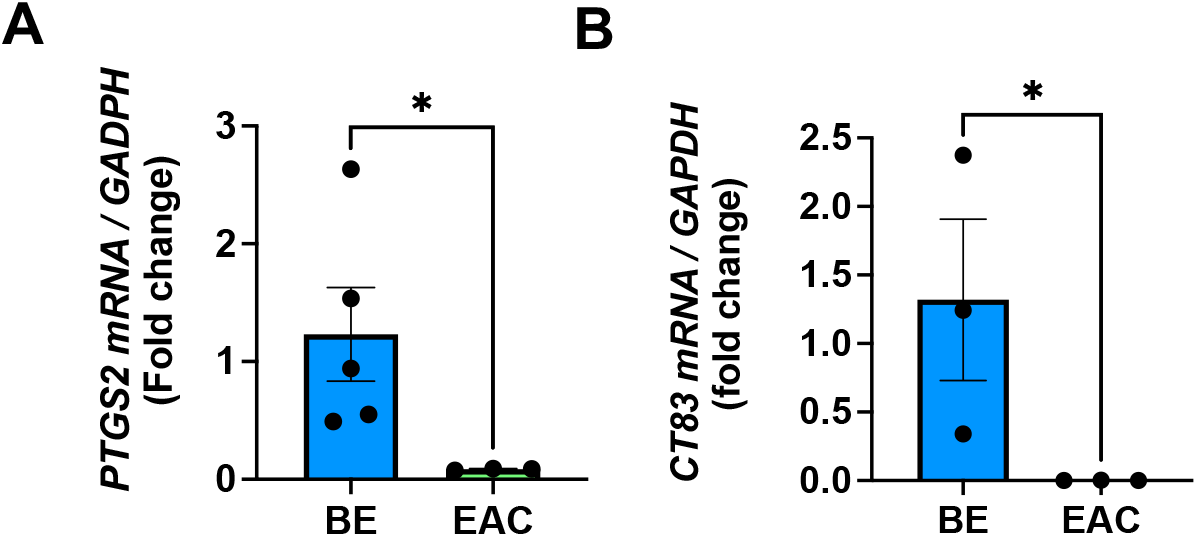
PTGS2 and CT83 are highly upregulated in BE compared with EAC. **A,B** Epithelial organoids generated from BE (n=3-5) or EAC (n=3-5) tissue were analyzed for **(A)** *PTGS2* and **(B)** *CT83* gene expression by real time PCR. (Data shown as mean +/− SEM, unpaired t test, significance: *, *p* <0.05).

**Expanded View Figure 2-.**
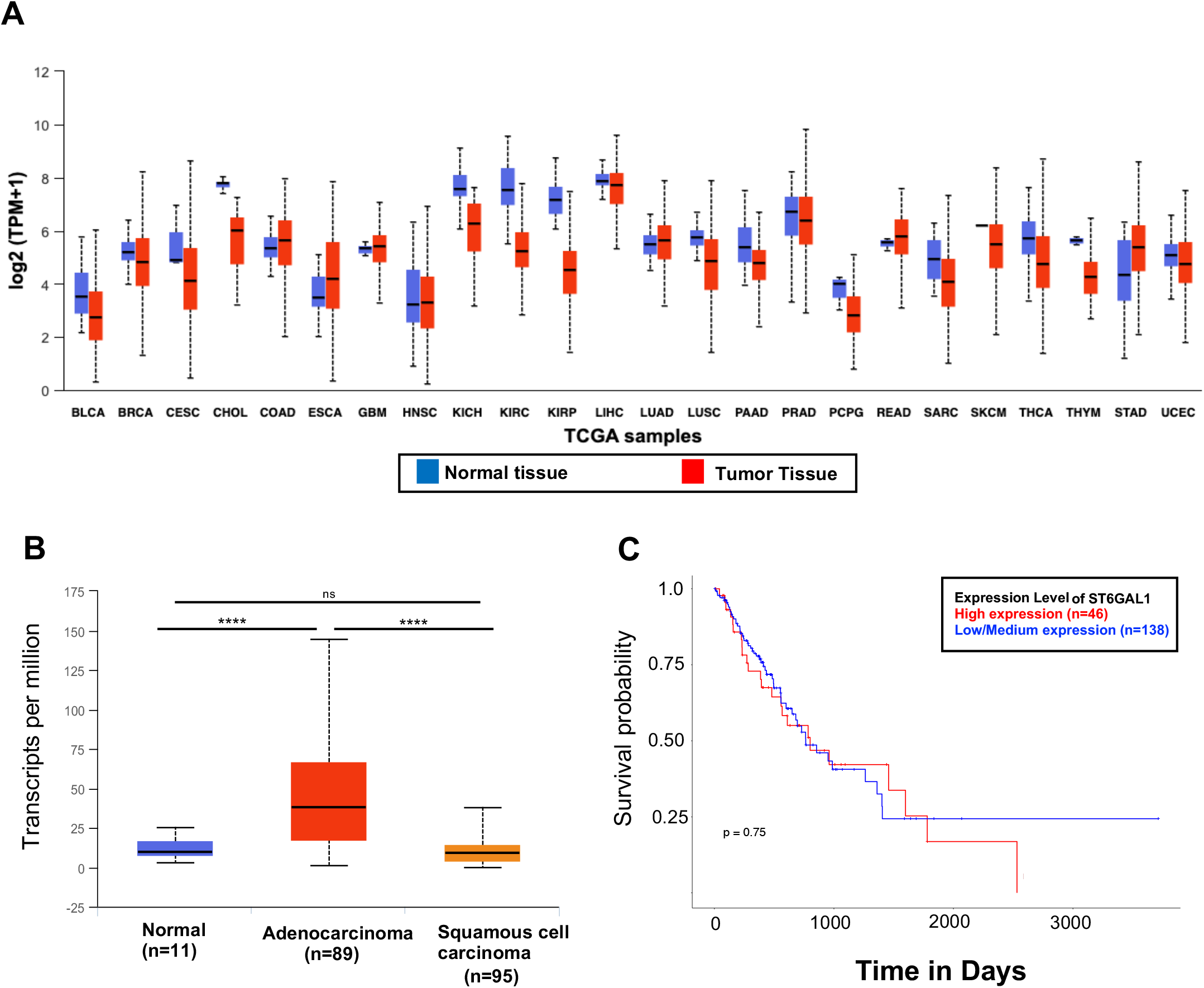
ST6GAL1 is highly expressed in EAC. **A-C** The UALCAN (The University of Alabama at Birmingham Cancer data analysis portal) online database was used to query TCGA (The Cancer Genome Atlas) public data. ST6GAL1 expression across 24 cancer types, including esophageal carcinoma (ESCA) and stomach adenocarcinoma (STAD). BLCA (bladder urothelial carcinoma), BRCA (breast invasive carcinoma), CESA (cervical squamous cell carcinoma), CHOL (cholangiocarcinoma), COAD (colon adenocarcinoma), ESCA, GBM (glioblastoma multiforme), HNSC (head and neck squamous cell carcinoma), KICH (kidney chromophobe), KIRC (kidney renal clear cell carcinoma), KIRP (kidney renal papillary carcinoma), LIHC, (liver hepatocellular carcinoma) LUAD (lung adenocarcinoma), LUSC (lung squamous cell carcinoma), PAAD (pancreatic adenocarcinoma), PRAD (prostate adenocarcinoma), PCPG (pheochromocytoma and paraganglioma), READ (rectal adenocarcinoma), SARC (sarcoma), SKCM (skin cutaneous melanoma), THCA (thyroid carcinoma), THYM (thymoma), STAD, UCEC (uterine corpus endometrial carcinoma). (**B)** ST6GAL1 expression in normal esophageal, EAC and ESCC tissue using TCGA public data. (Significance, ****, *p* <0.00001, statistics performed by UALCAN data analysis portal). (**C)** ST6GAL1 expression level correlation with esophageal cancer patient survival using TCGA public data.

**Expanded View Figure 3:**
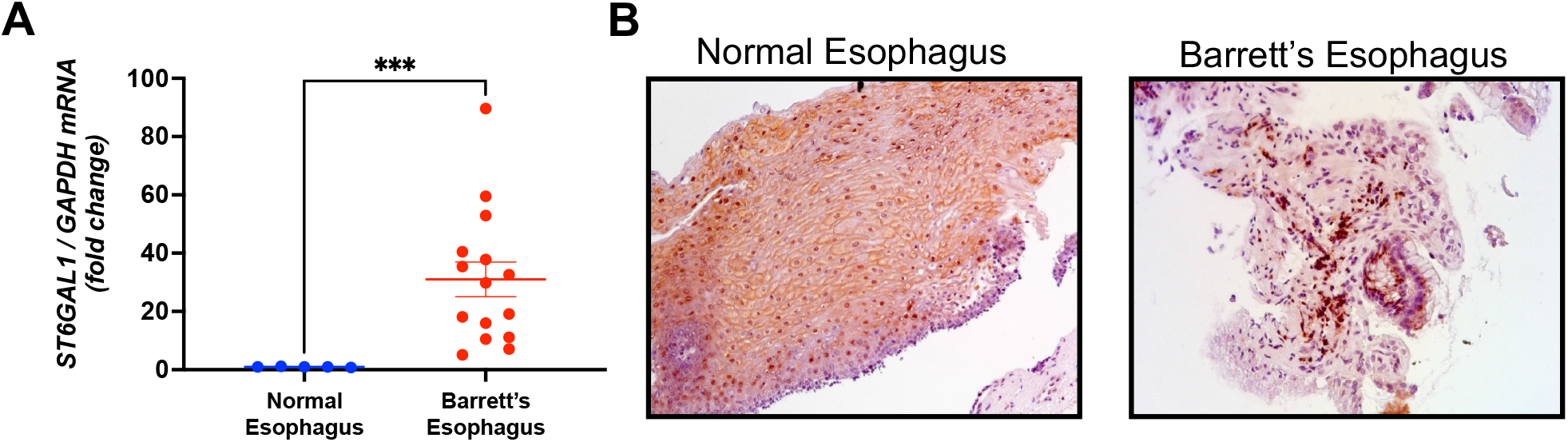
ST6GAL1 is not expressed in normal esophagus tissue. **A** Biopsies from normal esophagus (n=5) or BE (n=15) tissue were analyzed for *ST6GAL1* gene expression by real time PCR. (Data shown as mean +/− SEM, unpaired t test, significance: ***, *p* <0.0001). **B** Esophageal tissue from healthy donors or esophageal tissue from subjects with BE (LGD shown) stained for ST6GAL1 (DAB) by immunohistochemistry (n=3, representative images shown, 20×).

